# Genome mapping resolves structural variation within segmental duplications associated with microdeletion/microduplication syndromes

**DOI:** 10.1101/2020.04.30.071449

**Authors:** Yulia Mostovoy, Feyza Yilmaz, Stephen K. Chow, Catherine Chu, Chin Lin, Elizabeth A. Geiger, Naomi J. L. Meeks, Kathryn. C. Chatfield, Curtis R. Coughlin, Pui-Yan Kwok, Tamim H. Shaikh

**Affiliations:** Cardiovascular Research Institute, UCSF School of Medicine, San Francisco, California 94143, USA; Department of Integrative Biology, University of Colorado Denver, Denver, Colorado 80204, USA; Department of Pediatrics, Section of Clinical Genetics and Metabolism, University of Colorado School of Medicine, Aurora, Colorado 80045, USA; Department of Pediatrics, Section of Cardiology, University of Colorado School of Medicine, Aurora, Colorado 80045, USA; Department of Dermatology, UCSF School of Medicine, San Francisco, California 94143, USA; Institute for Human Genetics, UCSF School of Medicine, San Francisco, California 94143, USA

**Keywords:** Segmental duplications, genome mapping, structural variation, genomic disorders

## Abstract

Segmental duplications (SDs) are a class of long, repetitive DNA elements whose paralogs share a high level of sequence similarity with each other. SDs mediate chromosomal rearrangements that lead to structural variation in the general population as well as genomic disorders associated with multiple congenital anomalies, including the 7q11.23 (Williams-Beuren Syndrome, WBS), 15q13.3, and 16p12.2 microdeletion syndromes. These three genomic regions, and the SDs within them, have been previously analyzed in a small number of individuals. However, population-level studies have been lacking because most techniques used for analyzing these complex regions are both labor- and cost-intensive. In this study, we present a high-throughput technique to genotype complex structural variation using a single molecule, long-range optical mapping approach. We identified novel structural variants (SVs) at 7q11.23, 15q13.3 and 16p12.2 using optical mapping data from 154 phenotypically normal individuals from 26 populations comprising 5 super-populations. We detected several novel SVs for each locus, some of which had significantly different prevalence between populations. Additionally, we refined the microdeletion breakpoints located within complex SDs in two patients with WBS, one patient with 15q13.3, and one patient with 16p12.2 microdeletion syndromes. The population-level data presented here highlights the extreme diversity of large and complex SVs within SD-containing regions. The approach we outline will greatly facilitate the investigation of the role of inter-SD structural variation as a driver of chromosomal rearrangements and genomic disorders.

## Introduction

The sequencing and assembly of the draft human reference genome proved to be a tipping point in the field of human genetics; combined with the advent of affordable high-throughput short-read sequencing, it allowed the identification of variants in hundreds of thousands of individuals by aligning their sequencing reads directly to and comparing them with the reference genome. While genome sequencing has revealed the wide genetic diversity of human populations, short-read sequencing is unreliable for the analysis of areas of the human genome containing long, complex repetitive DNA elements, including telomeres, centromeres, and regions known as low copy repeats or segmental duplications (SDs). SDs are regions of ≥1000 bp that have two or more copies across the genome with ≥90% sequence identity (Lander et al. 2001; Bailey et al. 2001) and comprise ~5% of the human genome. SDs are often organized in blocks with complex internal structure, containing duplicated genomic segments, referred to as duplicons, that vary in order and orientation. A biologically significant subset of SDs are larger in size (tens or even hundreds of kb) with extremely high sequence identity (>99%) (Sharp et al. 2005), and are involved in recurrent genomic rearrangements (Lupski 1998; Bailey et al. 2001). Due to their length and sequence identity, these regions cannot be resolved using short-read sequencing, and in many cases, even with longer-read sequencing (Vollger et al. 2019).

The length and high sequence similarity of SDs make them an excellent substrate for non-allelic homologous recombination (NAHR), which occurs between highly identical paralogous copies of SDs and results in different types of structural variation (SV), including inversions, microdeletions, and microduplications (Stankiewicz and Lupski 2010; Carvalho and Lupski 2016). SDs are therefore hotspots of genomic rearrangements and complex SVs (Levy-Sakin et al. 2019), some of which give rise to genomic disorders caused by copy number variation (CNV) of dosage-sensitive or developmentally important genes (Lupski 2009, 1998; Bailey et al. 2001). Some of the well-studied examples include microdeletions at 7q11.23 known as Williams-Beuren syndrome (WBS, MIM #194050), 15q13.3 microdeletion syndrome (MIM #612001), 16p12.2 microdeletion syndrome (MIM #136570), and 22q11.2 Deletion Syndrome (MIM #188400), formerly known as DiGeorge syndrome (Peoples et al. 2000; Shaikh et al. 2000; Sharp et al. 2008; Girirajan et al. 2010). Microdeletion breakpoints in these syndromes are located in SDs, where NAHR between paralogous copies results in the deletion of genes within the interstitial single-copy region (Eichler 2001; Emanuel and Shaikh 2001; Carvalho and Lupski 2016).

Since SD-containing regions cannot be resolved using short-read sequencing technology, other techniques have been harnessed to reconstruct the structure of these regions. Some structural variation within SD-containing loci have been detected using low-resolution fluorescence *in situ* hybridization (FISH) analysis (Osborne et al. 2001; Sharp et al. 2008; Cuscó et al. 2008). Efforts to fully reconstruct SD-containing regions have required a scaffold of large-insert bacterial artificial chromosomes (BACs) in combination with long and short-read sequencing technologies (Huddleston et al. 2014; Steinberg et al. 2014), which would be both time- and cost-prohibitive for characterizing the SD-containing regions of a large number of samples. An approach to assemble SDs using only high-throughput long-read sequencing had trouble reconstructing paralogs longer than ~50 kb (Vollger et al. 2019).

Despite accumulating evidence for the role of SDs in disease-causing genomic rearrangements, their highly identical sequences, large size, and complex structures have made it difficult to elucidate their content. In this study, we demonstrate the use of the single molecule optical mapping technique to detect SVs over a wide range of sizes at the SD-containing regions of 7q11.23, 15q13.3 and 16p12.2. We present a suite of tools, OMGenSV, to use optical map data to genotype structural variation at any locus of interest, including complex genomic regions such as SDs. The high-throughput nature of the technique allowed us to analyze a diverse control dataset of 154 individuals; thus, we were able to determine the prevalence of SD configurations across different populations, including Africans, Americans, Europeans, and East and South Asians. Additionally, we applied our approach to patient samples with 7q11.23, 15q13.3 and 16p12.2 microdeletions and were able to further refine the microdeletion breakpoints. The ability to probe the internal structure of complex regions in a high-throughput manner and at low cost, using the techniques presented here, will greatly aid the study of structural variation within SDs and will help elucidate its relationship to chromosomal rearrangements, both in the general population and in individuals with genomic disorders.

## Results

### Optical mapping to elucidate the structural complexity within segmental duplications

Segmental duplications (SDs) are hotspots for rearrangements and large-scale structural variations (Emanuel and Shaikh 2001; Stankiewicz and Lupski 2002; Shaw et al. 2004). However, due to their complex structures and the high sequence identity shared by paralogous copies, SDs have been difficult to reliably map and sequence. We developed an approach that uses optical genome maps to identify the spectrum of structural configurations within and around SDs, and then to genotype those configurations in a high-throughput manner in a large set of samples. We were able to reliably map SD-containing regions at 7q11.23, 15q13.3, and 16p12.2 – all regions that are involved in recurrent structural variations associated with genomic disorders (Peoples et al. 2000; Sharp et al. 2008; Girirajan et al. 2010). We characterized these regions by optical mapping of 154 phenotypically normal individuals from 26 different populations comprising five super-populations: African (AFR), American (AMR), East Asian (EAS), European (EUR), and Southeast Asian (SAS) (Levy-Sakin et al. 2019).

*De novo* assembly of genome maps at these SD regions often resulted in assembly errors at paralogs with long stretches of identical label patterns. Thus, rather than using assembled contigs, our approach to genotyping SDs relied on identifying single molecules that mapped to a given SD configuration with labels anchored in unique regions on one or both sides (Fig. 1A). First, for each locus of interest, we compiled a list of potential configurations by examining all assembled local contigs (Fig. 1A (i)). Then, to evaluate the accuracy and prevalence of these configurations in our dataset, we aligned local single molecules from each sample to the set of potential configurations at that locus, filtering to retain molecules that aligned uniquely to one of the configurations (Fig. 1A (ii)) and whose alignment spanned both the identical region as well as its flanking context (Fig. 1A (iii)). The resulting set of aligned single molecules revealed the configuration(s) present in each sample (see Methods for additional details).

**Figure 1.**
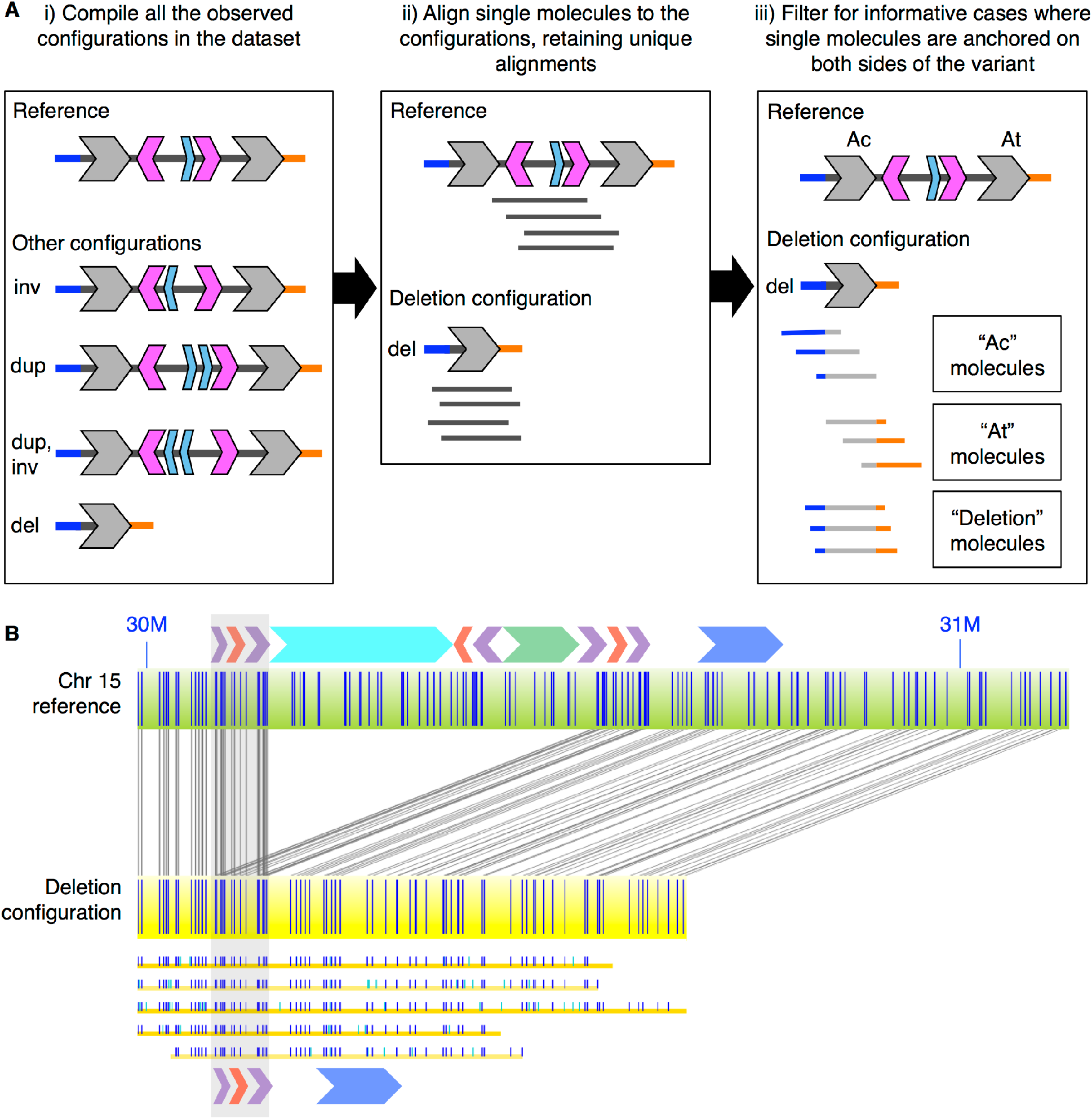
Optical mapping to genotype complex structural variation. A) Cartoon example of the pipeline (i-iii). (i), compilation of distinct configurations from all the assembled contigs in the full dataset. The cartoon locus depicted here includes inversions (inv), a duplication (dup), and a deletion (del). (ii) alignment of single molecules from each sample to the full set of local configurations seen in (i) to determine genotype. The example shown here has single molecule support for the reference and deletion (del) configurations. (iii) selection of informative molecules anchored in unique sequence flanking the repeat element. In the example shown here, Ac (A-centromeric) and At (A-telomeric) are the two paralogs of the duplicon marked by the grey arrow. The molecules labeled “Ac” and “At” cannot distinguish between the deletion and other configurations as they lack the full flanking context. The molecules labeled “Deletion” exclusively support the deletion configuration as they contain flanking sequences on both sides of the repeat element. B) A real example showing a deletion configuration at 15q13.3. The top green bar represents the reference configuration, while the middle yellow bar represents the deletion configuration. The SD duplicon structure corresponding to the reference are shown above the reference configuration as colored arrows. The yellow lines with blue and cyan vertical ticks below the deletion configuration are single molecules spanning the deletion. The SD duplicon structure corresponding to the deletion configuration are shown at the bottom. The grey transparent column highlights the breakpoint region that molecules needed to span in order to support the deletion.

### Structural variation at 7q11.23

7q11.23 contains two SDs – designated SD7-I and SD7-II – flanking a 1.3 Mb gene-containing region that is deleted in patients with WBS (Bayés et al. 2003). Known SVs within 7q11.23 include an inversion between SD7-I and -II, which was associated with a predisposition to the pathogenic deletion (Osborne et al. 2001), as well as deletions and duplications within the SDs (Cuscó et al. 2008).

We examined all assembled contigs from the 7q11.23 locus in our dataset of 154 diverse individuals, which revealed three major SVs in this region: the large inversion between the two SD7 regions (Fig. 2B), an inversion inside SD7-II (Fig. 2C), and a CNV within the A-centromeric (Ac) and A-telomeric (At) duplicons in SD7-I and SD7-II, respectively (Fig. 2D), here referred to as the A-CNV. The large inversion had breakpoints within the ~400 kb C-A-B duplicon block that was present in both SD7s (Fig. 2B). We found the inversion in 5% of observed configurations in the full dataset (3/62). We also detected a ~200 kb inversion of the A-middle (Am) duplicon within SD7-II (Fig. 2C), with an overall prevalence of 9% and a range of 4-20% within different populations.

**Figure 2.**
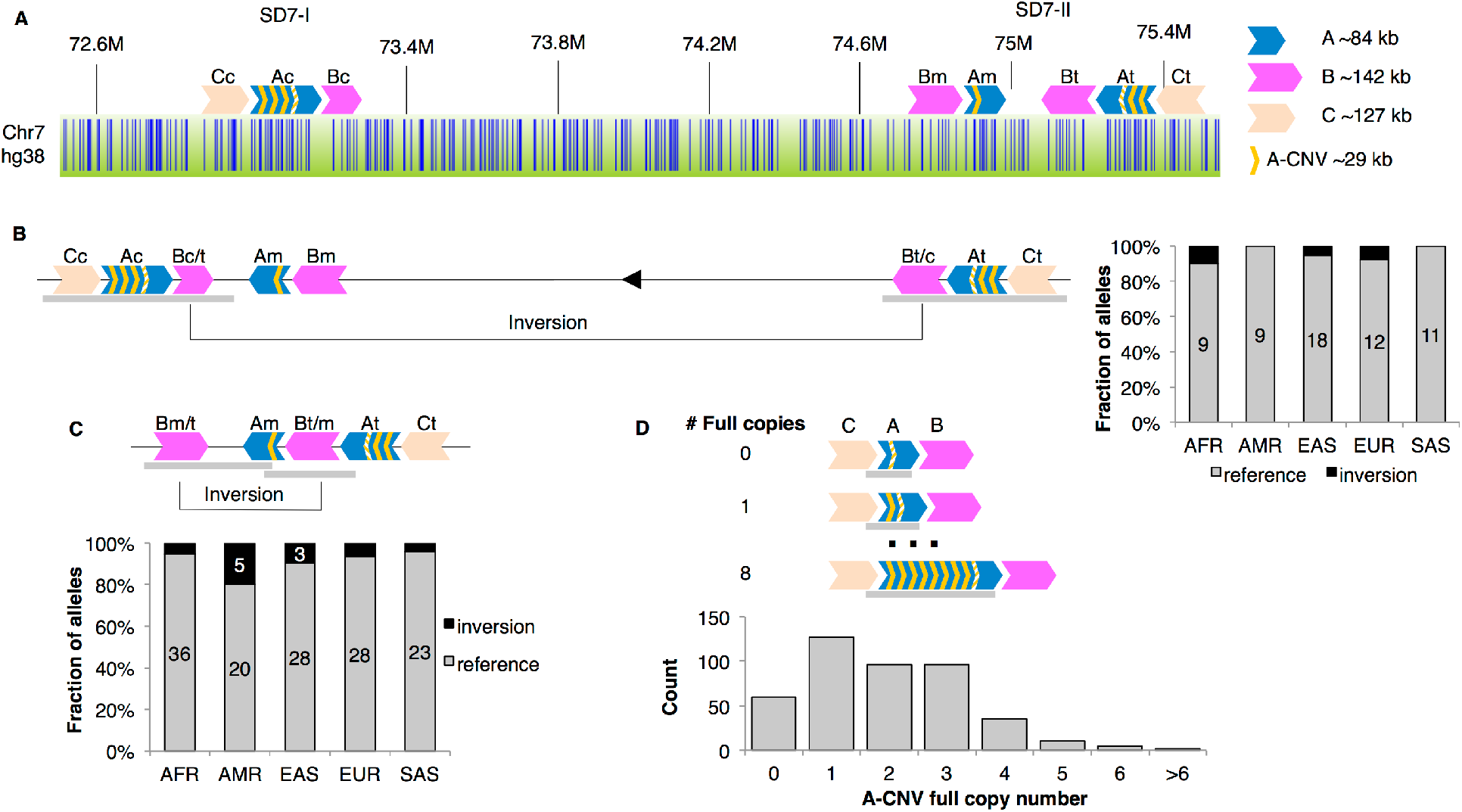
Structural variants at 7q11.23. A) The hg38 reference configuration of 7q11.23, showing duplicon positions and orientations for SD7-I and SD7-II. Paralogs are shown in the same color and are labeled e.g. ‘Ac’ and ‘At’ for the centromeric and telomeric copies of duplicon A. A partial copy of the A-CNV is marked with parallel lines. Below the duplicons, the optical map of this region is shown as a green bar with BspQI labels shown in blue. B) A large inversion observed between SD7-I and SD7-II, with breakpoints within the ‘C-A-B’ duplicon block. Right, a stacked bar graph showing the number of individuals carrying the large inversion allele or the reference configuration in each of the five populations covered in this study. C) A small inversion observed between Bm and Bt in SD7-II. Bottom, a stacked bar graph showing the number of individuals carrying the small inversion or the reference configuration in each of the five populations. For B) and C), labels on the bars show the number of times a configuration was detected in each population; count labels of one or two are not shown. D) A copy number variant observed in the A duplicon (A-CNV) flanked by the C and B duplicons in both SD7-I and SD7-II. Bottom, a bar plot depicting the full copy number alleles found in the A-CNV. No significant population differences were observed for (B), (C), or (D). In all SV diagrams, grey bars below the duplicons represent the critical regions that molecules needed to span in order to be informative for the configuration.

The A-CNV in duplicons Ac and At consisted of zero to eight full copies of a ~29-kb region, in addition to a consistent partial copy (Fig. 2D). In reference assembly GRCh38 (hg38), there were three full copies of this region in duplicon Ac and two full copies in At. We assessed the Ac and At CNVs together since they were embedded within the large C-A-B block in both SD7s and both paralogs were therefore flanked by long stretches of identical label patterns. The prevalence of each full copy number variant is presented in Fig. 2D: one copy was the most common, while copy numbers of >6 were rarely seen. Particularly long configurations can be difficult to detect if they require molecules to span regions substantially longer than the average molecule length (~250 kb), but the A-CNV copy numbers below 6x were unlikely to be adversely affected since they were smaller than the average molecule length (ranging from 94-235 kb for copy numbers 0-5). The A-CNV was highly variable both within and between individuals; 51% of samples had three different alleles at this CNV, while 25% had two alleles and 24% had four alleles, the latter group representing cases where Ac and At were both heterozygous for the A-CNV and each of the four alleles were distinct (Supplemental Fig. S1). Remarkably, none of the samples we analyzed had fewer than two distinct alleles. We did not detect any significant population-based differences in full copy number (Supplemental Fig. S2).

We also observed two other classes of configurations at the A-CNV locus (Supplemental Fig. S3). In one class, full copies of the CNV were interspersed with one or more partial copies; the most common such configuration is shown in Supplemental Figure S3A. Such partial copies were seen in 19 alleles from 17 individuals, all of African descent (Supplemental Fig. S3C), a significant enrichment over the other populations (*p* < 0.005 in each case, pairwise Fisher’s exact tests with Benjamini-Hochberg multiple testing correction). The other variant class involved a different label pattern immediately downstream of the CNV, shown in its most common configuration in Supplemental Figure S3B; this variant was present in nine samples of African, East Asian, and South Asian descent (Supplemental Fig. S3C) and was accompanied in different samples by 0-4 copies of the CNV. The number of distinct A-CNV alleles per sample (Supplemental Fig. S1) includes these variant alleles as well as the full copy number variants.

### Structural variation at 15q13.3

We next analyzed a region within 15q13.3 consisting of two segmental duplications, designated SD15-I and SD15-II, separated by a ~1.3 Mb unique region that is deleted in patients with the 15q13.3 microdeletion syndrome (Fig. 3A). This region is located in a highly unstable area of chromosome 15 that also includes the breakpoints for deletions associated with Angelman and Prader-Willi Syndromes (Amos-Landgraf et al. 1999; Christian et al. 1999).

**Figure 3.**
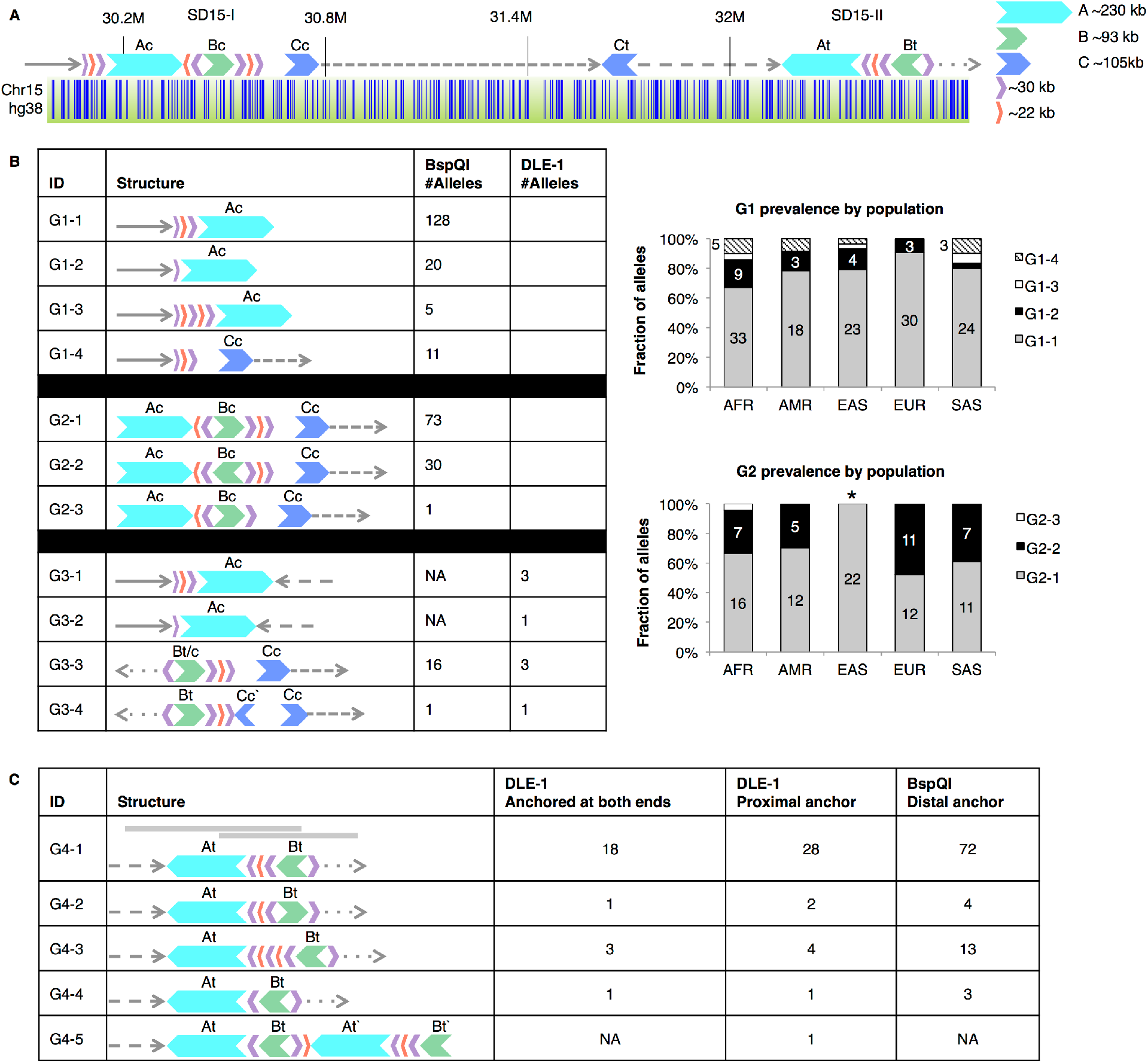
Structural variants at 15q13.3. A) The hg38 reference configuration of 15q13.3, showing duplicon positions and orientations for SD15-I and SD15-II. Paralogs are shown in the same color and are labeled e.g. ‘Ac’ and ‘At’ for the centromeric and telomeric copies of duplicon A. Grey arrows with different patterns mark the different unique regions flanking the SDs. Below the duplicons, the optical map of this region is shown as a green bar with BspQI labels shown in blue. B) Configurations anchored in the unique region either proximal or distal to SD15-I. Configurations were genotyped in three groups, G1, G2, and G3, using datasets labeled with the DLE-1 or the BspQI enzyme. For each genotyped sample, supporting molecules needed to span all of the duplicons and flanking unique regions depicted in the “structure” column. Right, stacked bar graphs showing the prevalence of configurations in the G1 (top) and G2 (bottom) groups for each of the five populations used in this study. Configuration G2-2 was significantly depleted in the East Asian population compared to all other populations (*p* < 0.05, pairwise Fisher’s exact test with Benjamini-Hochberg multiple testing correction comparing G2-1 and G2-2). Labels on the bars show the number of times a configuration was detected in each population. Count labels of one or two are not shown. C) Configurations anchored in the unique region either proximal or distal to SD15-II. The DLE-1 dataset contained molecules anchored within unique regions on both ends of SD15-II as well as molecules anchored only in the unique region proximal to SD15-II, while the BspQI dataset contained molecules anchored only in the unique region distal to SD15-II. Configurations were genotyped in one group, G4. Grey bars (top) indicate the proximal and distal critical regions: proximally anchored molecules extended at least to Bt, while distally anchored molecules extended at least to At. For (B) and (C), columns show the configuration IDs, their structure, and the number of alleles identified in our dataset with the indicated enzyme.

Within our dataset of 154 diverse individuals, we detected a large number of SVs at this locus, which we analyzed in groups based on the regions they shared in common, i.e. configurations anchored in the unique regions upstream or downstream of SD15-I and SD15-II (Fig. 3B,C). A subset of configurations represented inversions between the two SD15s, and their analysis required that supporting molecules be anchored in both upstream and downstream unique regions (Fig. 3B, group G3). One complicating factor in the analysis of this locus was the presence of a fragile site within the A duplicon for nicking enzyme BspQI; these fragile sites occur when two single-strand nicking sites occur close together on opposite strands, causing breakage of nicked DNA molecules so that few to no molecules are able to traverse the site. To overcome this limitation, we supplemented our dataset of 154 samples labeled using the BspQI nickase enzyme with a dataset containing 52 of those same samples labeled using the newer DLE-1 enzyme (Wong et al. 2020). DLE-1 deposits an epigenetic fluorescent label rather than nicking the DNA, and therefore does not create any fragile sites (Maggiolini et al. 2019). The 52 samples labeled with DLE-1 were selected from all 26 of the sub-populations in the original 154-sample dataset, with each sub-population represented by two samples.

From our analysis of SD15-I, we detected four different configurations anchored in the proximal unique region (Fig. 3B, G1), not including the inversions between SD15-I and SD15-II. The reference configuration (G1-1) was the most common (124/160 or 78% of observed configurations). Two additional configurations involved a contraction (G1-2, with a prevalence of 13%) or expansion (G1-3, prevalence of 3%) of the purple-red-purple duplicon triplet. The purple duplicons--which contain the *GOLGA8* genes--have previously been identified as key to the structural instability of this locus (Antonacci et al. 2014). An additional configuration in the proximal region of SD15-I involved a deletion of both the A and B duplicons (G1-4) and was seen in 7% of observed configurations.

We also detected three configurations anchored in the distal unique region of SD15-I (Fig. 3B, G2). The reference configuration (G2-1) was again the most common, representing 60% of observed configurations, while 25% of configurations had an inversion of the Bc duplicon (G2-2) (Antonacci et al. 2014). This inversion had strikingly different prevalence among populations, as it was seen most often in European samples (48%) and never in any East Asian samples. The East Asian population was significantly depleted in this configuration with respect to every other population (*p* < 0.05, pairwise Fisher’s exact tests with Benjamini-Hochberg multiple testing correction). The third configuration at this locus involved a contraction of the purple-red-purple duplicons (G2-3) and was only seen once in our dataset.

Inversions between SD15-I and SD15-II were detected by looking for molecules that aligned to unique regions either proximal to both regions or distal to both regions (Fig. 3B, G3). In the smaller DLE-1 dataset, which was used to traverse the BspQI fragile site within duplicon A for configurations G3-1 and G3-2, we detected inversions eight times. Configurations G3-3 and G3-4 were able to be assayed using the larger BspQI dataset, which showed evidence of inversion on 14% of observed configurations.

Configurations at SD15-II were assayed at the proximal end using DLE-1 to span duplicon At, while configurations at the distal end could be assayed using the full BspQI dataset (Fig. 3C). In some cases, molecules from the DLE-1 dataset were also able to span the full configurations from end to end. As in SD15-I, the reference configuration (G4-1) was the most prevalent (78% of observed configurations). Other configurations included an inversion of Bt (G4-2), analogous to but less frequent than the inversion of Bc in SD15-I (G2-2), and expansions and contractions of the purple-red-purple duplicon triplet (G4-3, G4-4). One chromosome harbored two tandem copies of At and Bt (G4-5), which was reconstructed from several tiled molecules from that sample because no single molecule spanned the full length of the configuration.

### Structural Variation at 16p12.2

We analyzed a region of 16p12.2 consisting of three segmental duplications, designated SD16-I, SD16-II, and SD16-III (Fig. 4A). The region between SD16-II and SD16-III is deleted in patients with 16p12.2 microdeletion syndrome (Ballif et al. 2007; Girirajan et al. 2010; Antonacci et al. 2010). Among the dataset of 154 diverse individuals, in addition to detecting haplotypes, corresponding to ‘S1’ and ‘S2’, reported previously (Antonacci et al. 2010) (Fig. 4B,C), we also detected novel SVs and configurations in this region. Most notably, we detected a novel inversion, ‘S3’, that combined the proximal part of S2, up through Bc`, with the distal part of the reference assembly (Fig. 4D). These long haplotypes were validated by tiling long single molecules (Supplemental Fig. S4).

**Figure 4.**
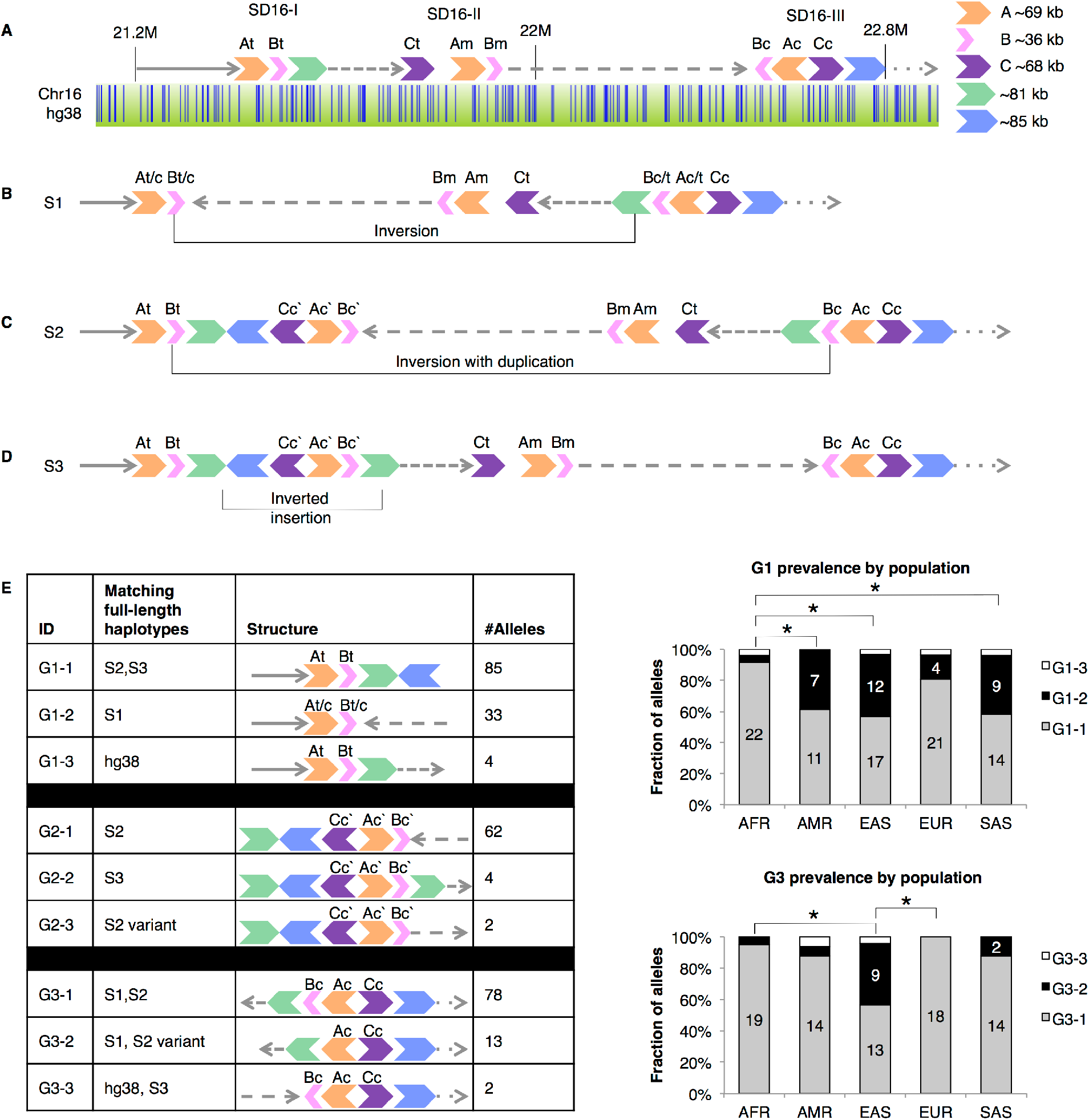
Structural variants at 16p12.2. A) The hg38 reference configuration of 16p12.2, showing duplicon positions and orientations for SD16-I, SD16-II, and SD16-III. Paralogs are shown in the same color, and are labeled e.g. ‘At,’ ‘Am,’ and ‘Ac’ for the telomeric, middle, and centromeric copies of duplicon A. Grey arrows with different patterns mark the different unique regions flanking the SDs. Below the duplicons, the optical map of this region is shown as a green bar with BspQI labels shown in blue. B) A large balanced inversion, “S1,” between At and Ac. C) A large inversion with duplication, “S2”. Newly created duplicons are marked as e.g. Cc`. D) A small inverted insertion detected distal to SD16-A on the reference configuration, labeled “S3.” E) Left, configurations genotyped in three groups, G1, G2, and G3, are shown. G1 configurations are anchored in the unique region proximal to SD16-I. G2 configurations are anchored in the green-blue duplicon pair that was not seen in the reference configuration. G3 configurations are anchored in the unique region distal to SD16-III. Columns depict the configuration ID, the longer haplotypes with which they are consistent (among the hg38 reference, S1, S2, and S3), the structure, and the number of alleles detected in the BspQI dataset. Supporting molecules for a given configuration had to span each of the depicted duplicons. Right, stacked bar graphs showing the prevalence of configurations from group G1 (top) and G3 (bottom) across the five populations included in this study. * *p* < 0.05, pairwise Fisher’s exact test with Benjamini-Hochberg multiple testing correction using the two most prevalent configurations in the group. Labels on the bars show the number of times a configuration was detected in each population. Count labels of 1 were not shown.

To genotype the variants in the full dataset, we grouped configurations based on shared location, as in 15q (Fig. 4E). Group 1 consisted of three configurations anchored in the unique region upstream of SD16-I (Fig. 4E, G1). G1-1 was the most common configuration, seen in 85/122 of observed configurations (70%). Between the two dominant configurations (G1-1 and G1-2), there was a significant difference in prevalence between the AFR population and the AMR, EAS, and SAS populations, with G1-1 comprising 92% of AFR G1 alleles but only 57-61% of G1 alleles from the other three groups (*p* < 0.05, Fisher’s exact test). Notably, a previous study of 24 chromosomes (Antonacci et al. 2010) found no support for the reference genome configuration at this locus. Our optical mapping results of 154 individuals showed that the hg38 reference assembly structure at this locus (G1-3) does exist in the general population, albeit at low frequency (3.2%, Fig. 4E, Supplemental Fig. S4A).

Group 2 consisted of three configurations, each corresponding to the extension of SD16-I seen in several of the rearranged haplotypes (Fig. 4C,D) but not in the reference. Of these, G2-1 was the most common configuration, found in 62/68 alleles (91%). Group 3 consisted of four configurations that were all anchored in the unique region downstream of SD16-III (Fig. 4E). The most common configuration was G3-1, with 78/93 G3 alleles (84%). Among the two most common G3 configurations, G3-2 was significantly enriched among the EAS population compared to AFR and EUR, comprising 39% of EAS G3 alleles but only 5% and 0% of AFR and EUR G3 alleles, respectively (p < 0.05, Fisher’s exact test).

### Breakpoint Mapping in Microdeletion Patients

We generated optical maps in four patients with microdeletions at the three loci analyzed in this study, with the goal of refining the microdeletion breakpoints.

We analyzed two patients with WBS, caused by microdeletions in 7q11.23, and reconstructed the structure of the deletion alleles in both. We found two different microdeletion configurations of 7q11.23 in the patients (Fig. 5A,B). In the first patient, the breakpoints were localized to Ac in SD7-I and Am in SD7-II, while in the second, they were localized to Bc in SD7-I and Bm in SD7-II, consistent with previous reports showing variation in the breakpoints of 7q11.23 microdeletions (Bayés et al. 2003; Merla et al. 2010).

**Figure 5.**
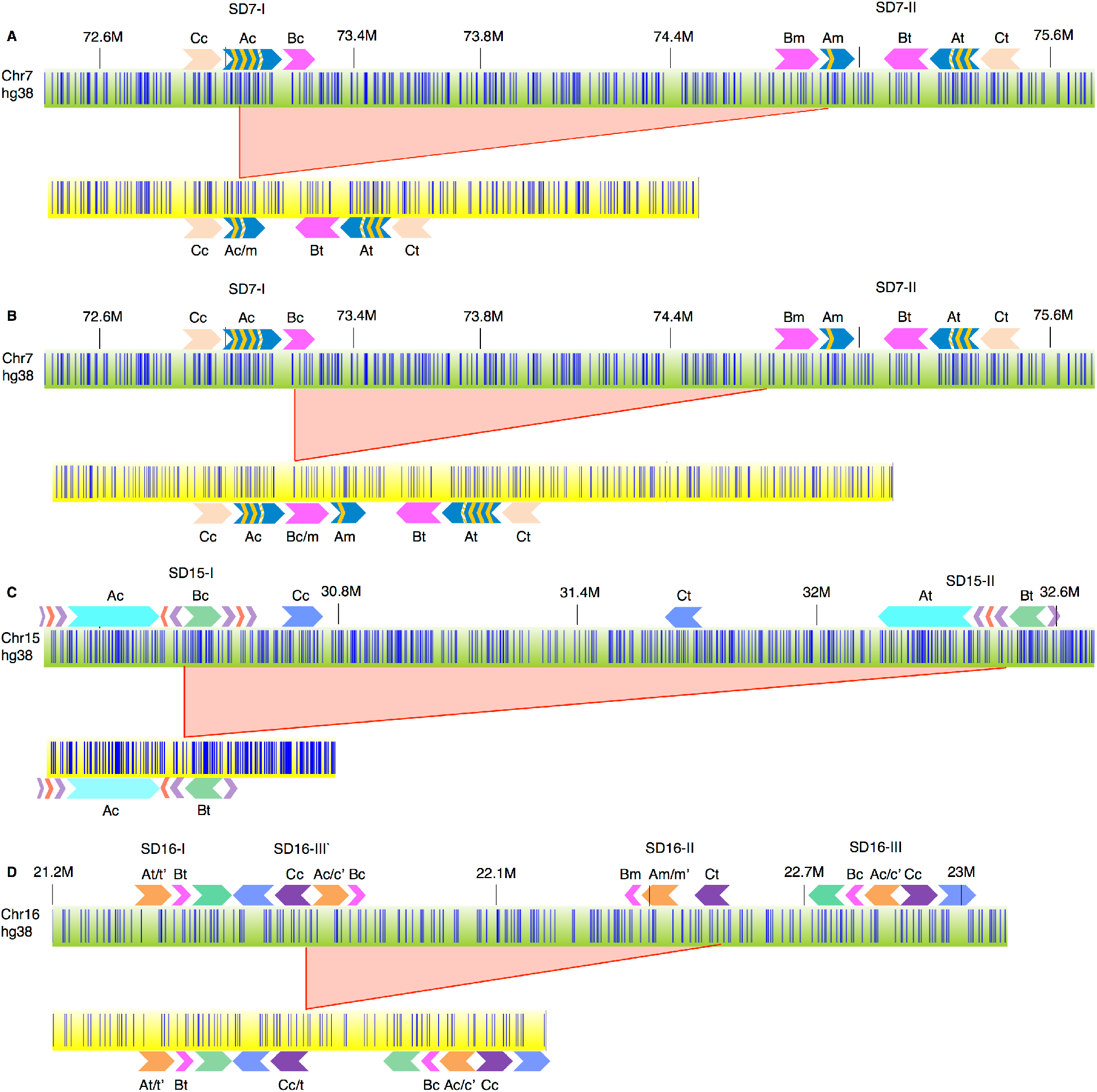
Breakpoint mapping in microdeletion patients. For each panel, the green bar (top) is the hg38 reference configuration of a given region, while the yellow bar (bottom) is the configuration of the deletion observed in the patient. Yellow bars in (A-C) depict assembled contigs while the yellow bar in (D) depicts a molecule. Red filled-in triangles indicate the region deleted in patients. Duplicon structures for the reference and deleted configurations are depicted above and below the bars, respectively. A) Patient 1 with 7q11.23 deletion. B) Patient 2 with 7q11.23 deletion. C) Patient with 15q13.3 deletion. D) Patient with 16p12.2 deletion.

Similarly, we reconstructed the deletion allele for one patient with the 15q13.3 microdeletion and one with the 16p12.2 microdeletion. In the 15q13.3 microdeletion, we localized the deletion breakpoints to the red and purple modules between Ac and Bc in SD15-I on one side and between At and Bt in SD15-II on the other side (Fig. 5C). In the 16p12.2 microdeletion patient, we found that the microdeletion breakpoints were localized to the C duplicons in SD16-II and SD16-III (Fig. 5D). Thus, optical mapping allowed us to reconstruct deletion haplotypes in patients with microdeletion syndromes and refine the microdeletion breakpoints within complex SDs.

## Discussion

Segmental duplications (SDs) are complex genomic structures containing long paralogs with extremely high sequence similarity, making them an excellent substrate for non-allelic homologous recombination (NAHR) (Stankiewicz and Lupski 2010). This process results in chromosomal rearrangements including microdeletions, microduplications, and inversions (Cuscó et al. 2008; Antonacci et al. 2010, 2014; Merla et al. 2010; Osborne et al. 2001; Hobart et al. 2010). Despite the importance of resolving the internal structure of SDs to elucidate the underlying cause(s) of associated chromosomal rearrangements, little is known about how these structures vary between individuals. Existing techniques are typically either unable to accurately assemble these regions due to short read lengths--with some repeat units extending for hundreds of kb, even "long-read" technologies such as PacBio or Nanopore are insufficient to reconstruct many SDs (Vollger et al. 2019)--or are too cost- and/or labor-intensive to be applied in a high-throughput study (e.g. "long-read" sequencing of individual BAC clones(Huddleston et al. 2014) or fiber FISH analysis (Molina et al. 2012)). Bionano optical mapping overcomes both of the above constraints by obtaining long read lengths with a fast and (at ~$600 per sample) cost-effective methodology.

Because automated *de novo* assembly of optical maps is error-prone at loci with long repetitive regions, we created software tools to rapidly genotype SVs in a high-throughput manner using unassembled single molecules. By applying this approach to a diverse dataset of 154 phenotypically normal individuals, we identified dozens of structural variants--some novel and some previously reported--at three different SD-containing loci associated with genomic disorders (7q11.23, 15q13.3, and 16p12.2) and created catalogs of SV patterns in these SD-containing loci, several of which had frequencies that varied by population. We also mapped the microdeletion breakpoints in patients with each of these genomic disorders. The resolution of the optical maps allowed us to narrow down SV breakpoints to individual duplicons inside SDs, which may help facilitate further breakpoint localization at single nucleotide-resolution.

Each of the loci in the study was found to contain at least one SV with significantly different frequency between populations (Fig. 3B, Fig. 4E, Supplemental Fig. S3). Importantly, two of these SVs affected the length of directly-oriented paralogous duplicons between SDs, which could serve as a template for NAHR to lead to pathogenic microdeletions and microduplications. At 15q13.3, a previously-reported inversion of the Bc module (Antonacci et al. 2014) had an overall prevalence of 29% (Fig. 3B, G2-2); it represented 48% of European alleles but was entirely absent in East Asian samples, similar to the population stratification of this SV observed in Antonacci et al. 2014 (Antonacci et al. 2014). This SV represents a substantial increase in the length of directly-oriented duplicons between the two blocks of SD and may therefore be predisposing to the pathogenic microdeletion/duplication (Antonacci et al. 2014), suggesting that East Asian populations would be expected to have low levels of 15q13.3 microdeletion/microduplication syndrome, while European populations may have the highest levels. However, Antonacci et al. 2014 suggested that the Bc inversion haplotype is not enriched among microdeletion patients and that the microdeletion mechanism may therefore not be strongly reliant on length of homology. A comprehensive analysis of SD configurations in parental samples may be required to evaluate the true impact of this SV on the microdeletion mechanism.

Similarly, at 16p12.2, configuration G1-2 (Fig. 4E) distinguished the S1 variant (Fig. 4B) from either the reference (Fig. 4A) or the S2 and S3 configurations (Fig. 4C,D). Notably, the S1 configuration lacks directly-oriented paralogs flanking the region that, on the reference, lies between SD16-II and SD16-III, which is deleted in the pathogenic microdeletion, while the other configurations have directly-oriented C duplicons flanking the region, suggesting that the S1 configuration may be protective against the pathogenic microdeletion (Antonacci et al. 2010). Accordingly, mapping of a 16p12.2 microdeletion patient sample showed a configuration consistent with a transmitting chromosome containing haplotype S2, with deletion breakpoints inside the C duplicons (Fig. 5D). The G1-2 configuration, representing the putatively protective S1 haplotype, varied in prevalence from 4% in Africans to 38-40% in East Asians, Americans, and South Asians, consistent with a previous study (Antonacci et al. 2010), suggesting that the latter three populations may be expected to have a lower prevalence of the pathogenic microdeletion. The third SV that varied in prevalence between populations was the partial A-CNV configuration at 7q11.23 (Supplemental Fig. S3), which was seen exclusively in African samples and would likely not affect predisposition towards the pathogenic microdeletion/microduplication.

While optical mapping of SD loci is a major advance, it is limited by the physical lengths of the molecules. Even with mean molecule N50 of 262 kb (range 199-395 kb), we are unable to genotype SVs containing repetitive elements where the length of the repeat exceeds the lengths of the molecules. A clear example of how the distribution of molecule lengths affects different SVs can be seen at the 7q locus. Genotyping the large inversion compared to the reference configuration (Fig. 2B) required finding molecules that spanned the entire length of the C-A-B duplicons with several flanking labels on both sides, i.e. molecules that were at least ~410 kb long. Molecules that spanned this range were only detected in 62/154 samples. In contrast, the CNV inside the A duplicons (Fig. 2D) required molecules that ranged from only 94 kb to 264 kb, depending on the allele, and consequently we were able to genotype at least two alleles in all 154 samples, with the vast majority of samples having at least three alleles between the two paralogs (Supplemental Fig. S1). Generally speaking, configurations that required molecules from the tail end of the length distribution for genotyping were less likely to be genotyped in any given sample. Future methodological refinements to increase molecule lengths will facilitate the genotyping of SDs with increasingly longer paralogous regions.

The molecular length is affected by the physical handling of long DNA molecules and experimental protocol. Our study was performed with the original labeling protocol based on a single-strand nicking enzyme that produces shorter molecules when two single-strand nicks are within several hundred basepairs on opposite strands, leading to fragile sites that break easily. The fragile sites not only shorten the molecular length, but also create breakpoints in the genome where no molecules can span across. This breakpoint problem is resolved with the use of a new labeling enzyme, DLE-1, which creates no fragile sites because it uses an epigenetic mark for labeling instead of single-strand nicks (Maggiolini et al. 2019).

This study demonstrates a high-throughput, cost-effective approach for characterizing SD-containing regions that involves using ultra-long optical maps to span the paralogous duplicons and capture the vast structural variability of these regions. Given our finding that SV patterns in the three SD-containing loci studied are highly variable and differ significantly between populations, including some that are likely to affect predisposition to pathogenic microdeletions/duplications, catalogs of SV patterns in these loci are most helpful in analyses of these regions in population and patient studies. Applying our high-throughput, cost-effective approach to additional complex loci throughout the genome will lead to catalogs of SV patterns that represent the genetic diversity of these regions and may reveal patterns that are prone to rearrangements that lead to genomic disorders. Furthermore, the tools and methods developed here will enable us to more accurately genotype SD configurations in clinical samples, which may consequently improve our ability to predict the risk for the occurrence of SD-mediated rearrangements associated with genomic disorders.

## Methods

### Human Subjects and Cell lines

Patient blood samples were obtained after informed consent, under the Institutional Review Board approved research protocol (COMIRB # 07-0386) at the University of Colorado Denver, School of Medicine.

### High-molecular-weight DNA extraction

High molecular-weight DNA for genome mapping was obtained from whole blood. White blood cells were isolated from whole blood samples using a ficoll-paque plus (GE healthcare) gradient. The buffy coat layer was transferred to a new tube and washed twice with Hank’s balanced salt solution (HBSS, Gibco Life Technologies). A small aliquot was removed to obtain a cell count before the second wash. The remaining cells were resuspended in RPMI (Gibco Life Technologies) containing 10% fetal bovine serum (Sigma) and 10% DMSO (Sigma). Cells were embedded in thin low-melting-point agarose plugs (CHEF Genomic DNA Plug Kit, Bio-Rad). Subsequent handling of the DNA followed protocols from Bionano Genomics using the Bionano Prep Blood and Cell Culture DNA Isolation Kit. The agarose plugs were incubated with proteinase K at 50°C overnight. The plugs were washed and then solubilized with GELase (Epicentre). The purified DNA was subjected to 45min of drop-dialysis, allowed to homogenize at room temperature overnight, and then quantified using a Qubit dsDNA BR Assay Kit (Molecular Probes/Life Technologies). DNA quality was assessed using pulsed-field gel electrophoresis.

### DNA labeling

7q11.23 proband DNA and 16p12.2 proband DNA was labeled using the IrysPrep Reagent Kit (Bionano Genomics). Specifically, 300 ng of purified genomic DNA was nicked with 10 U of nicking endonuclease Nt.BspQI [New England BioLabs (NEB)] at 37° for 2 hr in buffers BNG3 or BNG2, respectively. The nicked DNA was labeled with a fluorescent-dUTP nucleotide analog using Taq polymerase (NEB) for 1 hr at 72°. After labeling, the nicks were ligated with Taq ligase (NEB) in the presence of dNTPs. The backbone of fluorescently labeled DNA was counterstained with YOYO-1 (Invitrogen).

15q13.3 proband DNA was labeled using the Bionano Prep Early Access Direct Labeling and Staining (DLS) Kit (Bionano Genomics). 750 ng of purified genomic DNA was labeled by incubating with DL-Green dye and DLE-1 Enzyme in DLE-1 Buffer for 2 hr at 37°C, followed by heat-inactivation of the enzyme for 20 min at 70°C. The labeled DNA was treated with Proteinase K at 50°C for 1 hr, and excess DL-Green dye was removed by membrane adsorption. The DNA was stored at 4°C overnight to facilitate DNA homogenization and then quantified using a Qubit dsDNA HS Assay Kit (Molecular Probes/Life Technologies). The labeled DNA was stained with an intercalating dye and left to stand at room temperature for at least 2 hrs before loading onto a Bionano Chip.

### Data collection and assembly

The DNA was loaded onto the Bionano Genomics IrysChip for the 16p12.2 proband sample and the Saphyr Chip for all other probands and was linearized and visualized by the Irys or Saphyr systems, respectively. The DNA backbone length and locations of fluorescent labels along each molecule were detected using the Irys or Saphyr system software. Single-molecule maps were assembled *de novo* into genome maps using the assembly pipeline developed by Bionano Genomics with default settings (Cao et al. 2014).

### Cataloging and genotyping of structural variation at loci of interest

Structural variation at the 7q11.23, 15q13.3, and 16p12.2 loci was analyzed in a dataset of 154 phenotypically normal individuals from 26 diverse sub-populations, with genome maps labeled using the Nt.BspQI single-strand nickase enzyme (Levy-Sakin et al. 2019, NCBI BioProject PRJNA418343). For 15q13.3, one of the duplicons contained a fragile site wherein two Nt.BspQI label sites were in close proximity on opposite strands, resulting in consistent DNA breakage between the sites. For a complete analysis that could span that position, we included a dataset labeled with the DLE-1 enzyme, which uses an epigenetic label rather than nicking DNA and therefore does not produce fragile sites (Maggiolini et al. 2019). This DLE-1 dataset contained 52 samples from the original dataset, with two samples per sub-population (Wong et al. 2020, NCBI BioProject PRJNA611454).

Structural variation at the three loci was assessed using the Optical Maps to Genotype Structural Variation (OMGenSV) package as described in Demaerel et al. 2019 with minor modifications. To create a catalog of configurations for each locus, contig alignments from the exp_refineFinal1_SV folder of each sample’s de novo assembly were merged into a single alignment file, and contigs aligning to the loci of interest were visualized using the ‘anchor’ mode in OMView from the OMTools package (Leung et al. 2017). Distinct configurations were manually identified from this visualization, and corresponding CMAP files were made for each, including at least 500 Mb of unique flanking sequence where applicable. When SDs contained multiple SVs and were too long to analyze from end to end with single molecules (e.g. over ~350 kb), they were subdivided into groups that were typically anchored in the unique regions either upstream or downstream of the SD (Fig. 3B, Fig. 4E).

For each group of configurations, the corresponding CMAPs were compiled into a single file and used as input for the OMGenSV pipeline, along with local molecules from each sample and a set of ‘critical regions’ defining the area(s) on each CMAP that molecules need to span in order to support the presence of that configuration in the sample. OMGenSV then performs the following steps for each sample: 1) align local molecules to the configuration CMAPs; 2) filter for molecules that have a single best alignment among the different CMAPs, excluding those that align equally well to two or more CMAPS; 3) filter for molecules whose best alignment spans a pre-defined critical region; 4) report supported configurations for each sample, and 5) identify configurations with weak support in a given sample that would benefit from manual evaluation. The molecule support for these cases was manually evaluated using OMView (Leung et al. 2017). Rarely, we found new configurations during the manual evaluation phase that had not been represented in assembled contigs; in these cases, we created CMAPs for the new configurations and included them in a new run of the pipeline.

For the A-CNV at 7q11.23, the ‘partial’ alleles (Supplemental Fig. S3A) were too similar to full-copy alleles to disentangle using the standard OMGenSV pipeline, so we modified the protocol described above as follows. First, we ran the pipeline using a set of CMAPs that included the C-A-B duplicon block with 0,1, 2, or 3 full copies of the A-CNV, as well as several open-ended configurations that contained only duplicons C and A. One open-ended configuration had A containing 4 full A-CNV copies and then terminating, acting as a ‘sink’ for molecules with 4 or more full copies. The last two open-ended C-A configurations included partial copies of the A-CNV: full, full, partial, full, and full, full, partial, partial, full. After running the pipeline with these CMAPs, any molecules that aligned best to either of the configurations containing partial copies of the A-CNV were filtered out, creating a pool of local molecules that were likely to only contain full copies of the A-CNV. These filtered sets of local molecules were used as input for a new run of the pipeline, using CMAPs containing the full C-A-B duplicon block with 0-8 copies of the A-CNV. This run was processed in the standard way described above, and used to generate the results for full copies of the A-CNV (Fig. 2D, Supplemental Fig. S2). Molecules from the first run that aligned to the configurations with partial A-CNV copies were manually inspected to genotype those samples. Samples containing the ‘downstream variant’ A-CNV alleles (Supplemental Fig. S3B) were genotyped by running the pipeline with a CMAP containing two open-ended configurations beginning at the end of A and continuing through B, containing either the canonical end of A or the ‘downstream variant’ end of A. All molecules that aligned best to the ‘downstream variant’ configuration were manually evaluated to genotype that sample.

### Breakpoint Mapping in Microdeletion Patients

For each proband, assembled contigs that aligned to the locus of interest were manually inspected to identify the microdeletion configuration. Once identified, the underlying single molecules were examined in order to verify that the microdeletion breakpoint was well-supported. In the case of the 16p12.2 microdeletion proband, no assembled contigs contained a microdeletion configuration. Instead, we aligned local single molecules to the hg38 reference and identified those whose alignments supported a microdeletion breakpoint. We found several long single molecules that showed the same consistent breakpoint, one of which is shown in Fig. 5D.

## Data Access

The optical map data generated in this study have been submitted to the NCBI BioProject database (https://www.ncbi.nlm.nih.gov/bioproject/) under accession number PRJNA626024.

## Acknowledgements

This work was made possible by a grant, GM120772 to T.H.S and P-Y.K, from the National Institute of General Medical Sciences of the National Institutes of Health (NIH).

## Author Contributions

T.H.S and P-Y. K conceived the study and analysis using Bionano optical mapping. Y.M and F.Y performed the analysis and interpretation. C.R.C, N.J.L.M and K.C.C obtained informed consent from parents and patients during hospital follow-up and facilitated collection of blood samples. C.R.C co-ordinated study and subject enrollment. E.A.G., S.K.C., C.C., and C.L. carried out sample processing and experimentation. Y.M., F.Y., T.H.S. and P-Y. K drafted and critically revised the article. Final approval of the version to be published was given by Y.M., F.Y., S.K.C., C.C., C.L., E.A.G., N.J.L.M., K.C.C., C.R.C., P.Y.K., and T.H.S.

## Disclosure Declaration

The authors declare no competing interests.

